# Measurement of adhesion and traction of cells at high yield (MATCHY) reveals an energetic ratchet driving nephron condensation

**DOI:** 10.1101/2024.02.07.579368

**Authors:** Jiageng Liu, Louis S. Prahl, Aria Huang, Alex J. Hughes

## Abstract

Engineering of embryonic strategies for tissue-building has extraordinary promise for regenerative medicine. This has led to a resurgence in interest in the relationship between cell biophysical properties and morphological transitions. However, mapping gene or protein expression data to cell biophysical properties to physical morphogenesis remains challenging with current techniques. Here we present MATCHY (<u>m</u>ultiplexed <u>a</u>dhesion and traction of <u>c</u>ells at <u>h</u>igh <u>y</u>ield). MATCHY advances the multiplexing and throughput capabilities of existing traction force and cell-cell adhesion assays using microfabrication and an automated computation scheme with machine learning-driven cell segmentation. Both biophysical assays are coupled with serial downstream immunofluorescence to extract cell type/signaling state information. MATCHY is especially suited to complex primary tissue-, organoid-, or biopsy-derived cell mixtures since it does not rely on *a priori* knowledge of cell surface markers, cell sorting, or use of lineage-specific reporter animals. We first validate MATCHY on canine kidney epithelial cells engineered for RET tyrosine kinase expression and quantify a relationship between downstream signaling and cell traction. We go on to create a biophysical atlas of primary cells dissociated from the mouse embryonic kidney and use MATCHY to identify distinct biophysical states along the nephron differentiation trajectory. Our data complement expression-level knowledge of adhesion molecule changes that accompany nephron differentiation with quantitative biophysical information. These data reveal an ‘energetic ratchet’ that explains spatial nephron progenitor cell condensation from the niche as they differentiate, which we validate through agent-based computational simulation. MATCHY offers automated cell biophysical characterization at >10^4^-cell throughput, a highly enabling advance for fundamental studies and new synthetic tissue design strategies for regenerative medicine.

## Introduction

Cell collective mechanics are the proximate cause of tissue morphogenesis (1) - the tissue growth, shape change, and compartment interfacial geometry that determine normal and diseased organ function (**Fig 1A**). Interactions between cell tension, adhesion, proliferation, and migration sculpt many organs and feed back on cell behavior, for example in heart tube looping, craniofacial development, and condensation of hair follicle, feather, gut villus, and limb bud/digit structures (2–8). The formation of blood-filtering nephron structures in the developing kidney is an archetypal example of the relationship between radical mechanical and morphological transitions (9, 10). Cap mesenchyme cells (the nephron progenitors) lie at the tips of the developing ureteric bud epithelial tree (the future urinary collecting ducts), differentiating in response to Wnt (11, 12) and other biochemical cues from the ureteric bud and surrounding stroma (13–21). Nephron progenitors periodically condense into early nephrons by mesenchymal-to-epithelial transition (MET) as they differentiate, in parallel with a transition in cytoskeletal and adhesion molecule expression typical among other METs (10, 22–26). However, this ‘cell state/biochemical layer’ of understanding has not yet been paired with a commensurate ‘biophysical layer’ of cell mechanical changes that guide nephron self-organization. The lack of a complete mechanochemical picture of development remains a significant barrier to tissue construction by developmental mimicry, a nascent paradigm in tissue engineering for regenerative medicine (27).

**Fig. 1:**
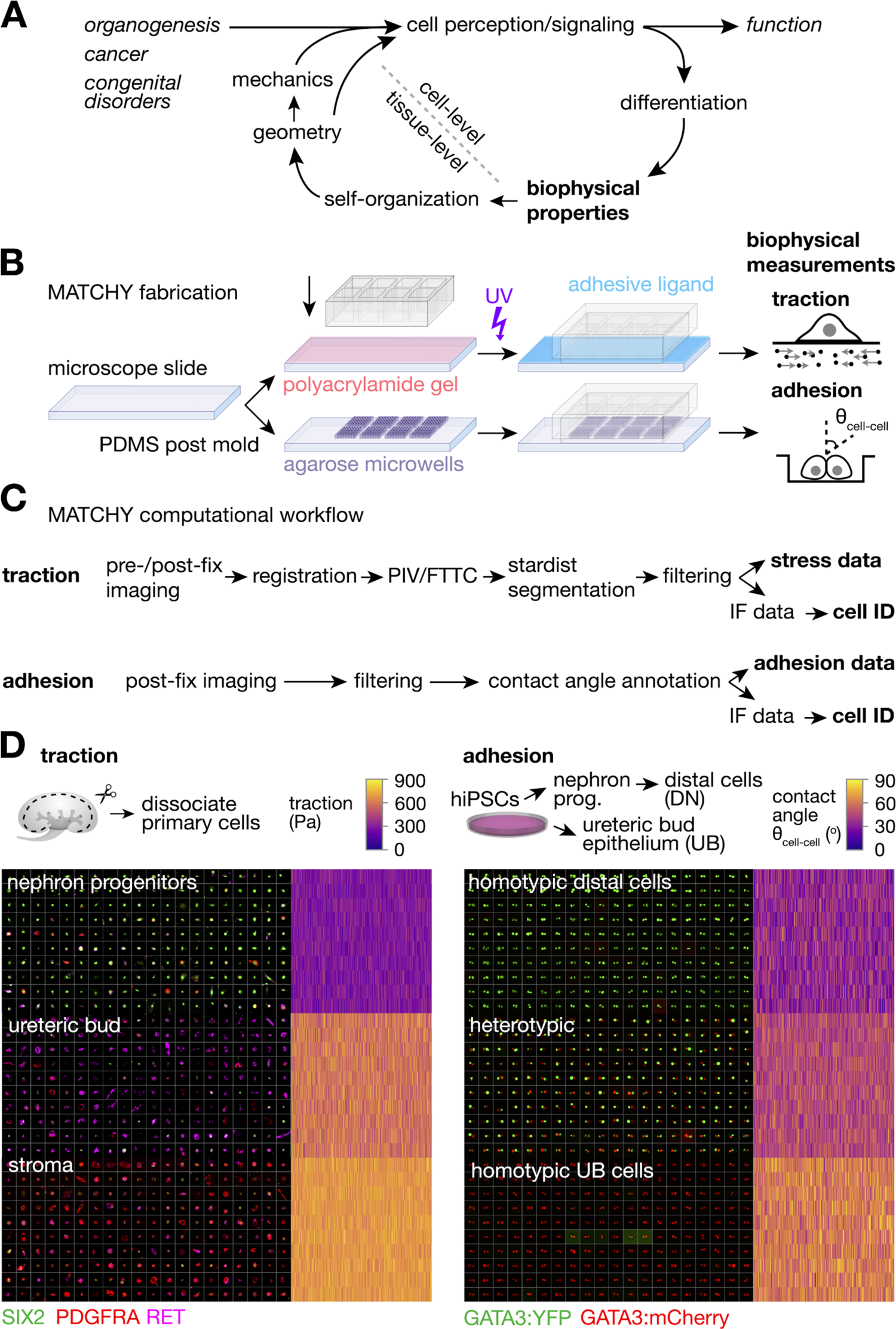
MATCHY quantifies biophysical parameters of cells in compositionally complex mixtures and associates them with cell type/signaling state at high throughput. (A) Schematic of biophysical contributions to tissue organization in development and disease. Cell transcriptional and signaling states manifest as biophysical properties that set tissue organization by migration, proliferation, and sorting. Cell traction and adhesion measurements therefore hold predictive value for tissue organization and function. (B) Schematic of microfabrication workflows for MATCHY traction and adhesion assays. (C) Schematic of automated computational pipeline for high-throughput cell biophysical quantitation and post-hoc assignment of cell type/signaling state from serial immunofluorescence analysis. (D) Example high-throughput traction and adhesion data for dissociated primary E17 mouse embryonic kidney cells and iPSC-derived kidney cell lineages respectively. Traction data is summarized as a mosaic image (*left*) of a subset of 200 cells per type category, along with a heatmap showing traction stress for 4,000 cells per category. Contact angle data is summarized similarly for a subset of 200 doublets of iPSC-derived kidney cell lineages per type category out of 2,000 doublets per category.

One difficulty that hobbles construction of biophysical-layer understanding across developmental systems is a lack of accessible tools for mechanical characterization of cells and tissues. Techniques such as atomic force microscopy (AFM) (28–30), micropipette aspiration (31–33), optical tweezers (34, 35), droplet deformation (36, 37), and traction-force microscopy (TFM) (38, 39) have yielded unprecedented advances here (40). However, each of these techniques suffers from low throughput, high technical complexity, or both. In parallel, researchers have inferred drivers of self-organization during morphogenesis from genetic model studies, primary cell self-organization assays, and cytoskeletal/adhesion expression profiling (33, 40–43). For example in nephron-forming niches, knockout of tension and adhesion regulators including non-muscle myosin II (Myh9/10) and integrin ɑ8β1 (ITGA8) alter niche and early nephron organization, affecting nephrogenesis rate (44, 45). Cells dissociated from embryonic kidneys spontaneously recover some aspects of native spatial structure, at least at short spatial length-scales. For example, ureteric bud aggregates (22) surrounded by SIX2+ nephron progenitors capable of rudimentary bona fide branching (46) and some connectivity with distal domains of nearby re-forming nephrons (47, 48). Researchers have made significant progress in defining a ‘cadherin code’ that distinguishes anatomical compartments, namely differential expression of cadherins between naive nephron progenitors (*Cdh2*, *4*, *6*) and those undergoing MET to renal vesicles and later stages associated with nephron segmentation (*Cdh1, 2, 3, 4, 6, 11, 16*) (22, 23, 25, 49, 50). However, cadherin expression alone is not necessarily predictive of self-organization outcomes (51). Though differential adhesion/interfacial tension has been raised as a compelling theory explaining niche and (more specifically) nephron self-organization (10, 22, 49, 51–54), it remains to be tested with direct biophysical measurements.

Here we present MATCHY (<u>m</u>ultiplexed <u>a</u>dhesion and traction of <u>c</u>ells at <u>h</u>igh <u>y</u>ield), which enables TFM and cell doublet adhesion assays to be integrated with immunofluorescence (IF)-based cell type and state measurement (**Fig. 1B-D**). MATCHY is powered by a computational pipeline designed for high-throughput and multiplexed measurements. We apply MATCHY to associate nephron progenitor lineages with biophysical states during mouse nephron development. We first validate MATCHY on a well-studied Madin-Darby canine kidney (MDCK) cell line engineered to express the receptor tyrosine kinase RET (55). RET functions through a ligand-receptor interaction with glial cell-derived neurotrophic factor (GDNF) to activate extracellular signal-related kinase (ERK) and other downstream signals to drive branching morphogenesis (56). ERK signaling, in turn, stimulates cell traction forces across epithelial tissue layers (57). We show that cell traction forces can be measured upon cell fixation rather than lysis, enabling downstream analyses such as immunofluorescence for phospho-ERK (active ERK) using a machine-learning toolbox for automated data analysis. We next apply MATCHY to measure traction forces produced by single cells in heterogeneous primary cell populations isolated from the mouse embryonic kidney ‘nephrogenic zone’. We leverage the ability of our pipeline to retrospectively link cell traction and identity in these heterogeneous cell mixtures. We then repeat a similar analysis for cell doublet adhesion assays in microwell arrays. These measurements together serve as a biophysical atlas of nephron progenitor lineage commitment, showing progressively increasing cell traction and homotypic adhesion along the nephron condensation ‘trajectory’. Heterotypic adhesion data reveal an ‘energetic ratchet’ that would tend to drive cells toward physical segregation from the nephrogenic niche, as observed *in vivo*. We show that biophysical data alone is sufficient to account for nephrogenic niche-like and early nephron structures using agent-based modeling and primary cell spheroid self-organization assays.

Together these data establish MATCHY as a flexible tool for mechanical analysis of cell mixtures and provide a biophysical basis for nephron formation by MET. By linking such data to organizational outcomes, we establish an engineering blueprint for synthetic nephrogenesis through cell engineering or other methods requiring initial or boundary biophysical conditions. Such data will inform future efforts to generate uniform, compact arrays of functional nephrons for kidney replacement duty. MATCHY is designed for application across a variety of cell systems, enabling biophysical characterization of organization across diverse applications in development, disease, and engineered tissues.

## Results

We designed MATCHY for simultaneous, multiplexed measurement of cell traction forces, cell-cell adhesion, and protein biomarker expression to correlate cell identities/states with biophysical information. For the traction arm of MATCHY, we fabricate polyacrylamide gel sheets with validated mechanical properties (58) bearing fluorescent microparticles as fiducial markers on standard microscopy slide substrates (**Fig. 1B**). Cell-adhesive ligands (specifically Matrigel in this study) are then photo-patterned onto the gel before assembling substrates into a modular culture well system. We reasoned that IF staining could be integrated with traditional traction force microscopy since cell fixation simultaneously relaxes cell tension (59) and permanently adheres cells to the polyacrylamide substrate. Indeed, reading out cell tension by fixation successfully recovered 72% of total traction magnitude measured by traditional cell lysis in a human induced pluripotent stem cell (iPSC) population (**Fig. S1**). We next created an automated experimental and image analysis process for high-throughput, ‘one-button’ image acquisition and pre-processing, multiplexed cell detection, traction force microscopy by fourier transform traction cytometry (FTTC) (60), and classification based on IF marker expression. This process enables high-throughput characterization of e.g. primary tissue- and iPSC-derived cell populations (**Fig. 1D**) (61). We validated the traction arm of MATCHY using a Madin-Darby canine kidney (MDCK) cell line overexpressing the human RET9 receptor (MDCK-RET9) (55). These cells change their adhesive properties and scatter in response to GDNF (55, 62) (**Fig. 2A**). Cells elevate pERK downstream of signaling complexes of RET and its co-receptor GFRA1 in response to GDNF (56, 63). Erk phosphorylation (pERK) stimulates contractile pulses within MDCK cell layers by activating the Rho/ROCK pathway (57). A similar action of RET through Rho has been observed in other cell types (64, 65). In 2D cell culture, we verified that addition of GDNF + GFRA1 induced MDCK-RET9 cell scattering, while untreated cells maintained intact colonies (**Fig. 2B**). However, a quantitative relationship between RET-GDNF signaling and traction force has not been determined. We quantified the traction force of MDCK-RET9 cells treated with GDNF + GFRA1 relative to negative controls consisting of untreated cells or those treated with the non-muscle myosin-II-specific ATPase inhibitor Y-27632 or myosin II inhibitor blebbistatin (**Fig. 2C-E**). GDNF + GFRA1 increased traction stresses by 1.6-fold, while inhibitor treatments significantly ablated traction stresses (**Fig. 2D**). Leveraging our ability to simultaneously measure RET marker expression and pERK level in the same cells, we noted a positive correlation between the two in the cell population for a fixed GDNF + GFRA1 stimulation (**Fig. 2E**). Traction stresses also monotonically increased with RET expression in the presence of GDNF + GFRA1 and indicated a possible switch-like increase in cell tension at intermediate expression. In untreated controls, there was no correlation between RET level and ERK phosphorylation or cell traction. These data indicate successful implementation of traction force microscopy integrated with multiplexed detection of protein markers of cell identity and state by immunofluorescence.

**Fig. 2:**
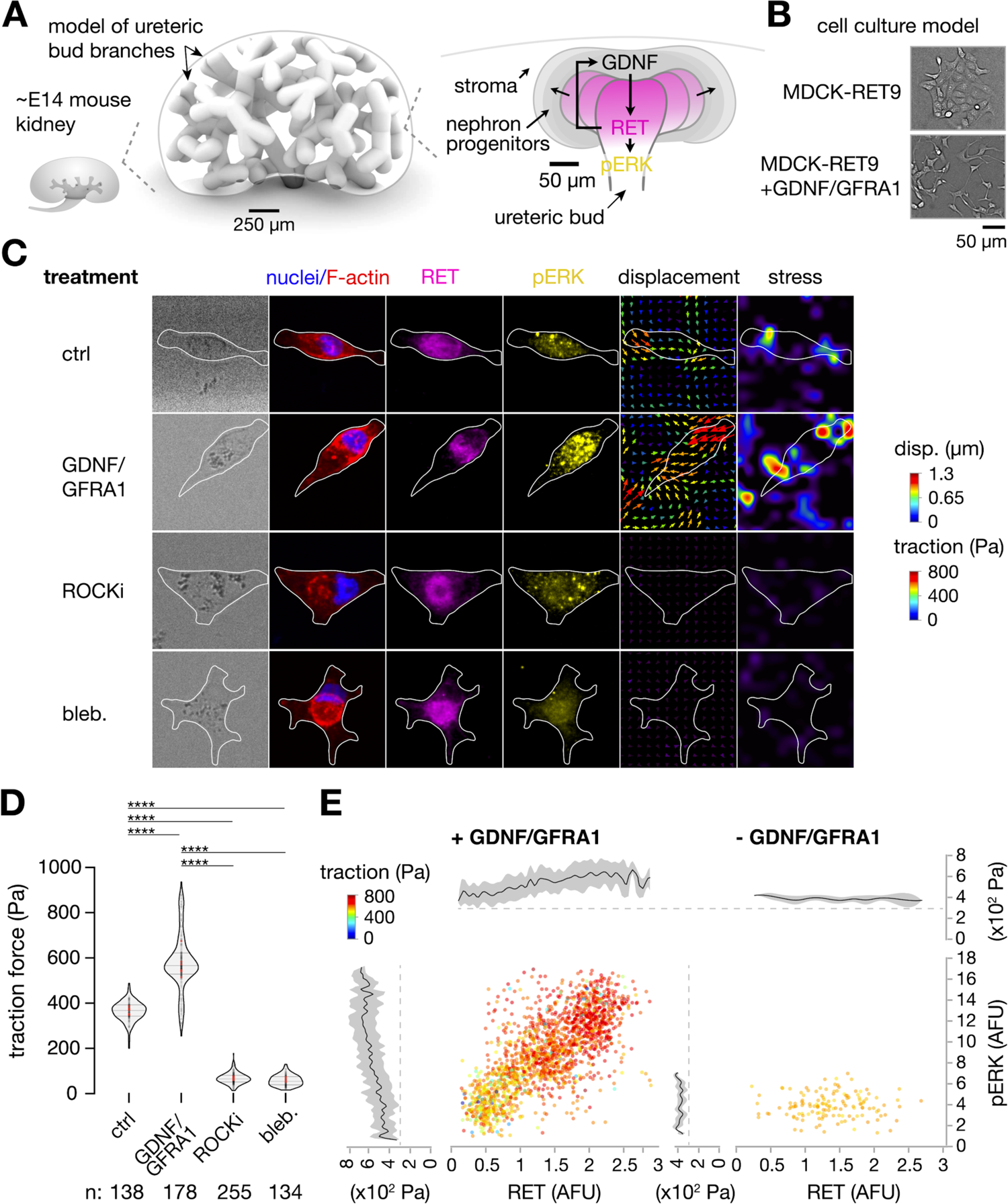
MATCHY captures cell traction effects of GDNF-RET signaling at high throughput. (**A**) Model schematic of ureteric bud tubule branching in the embryonic kidney. Right, detail of nephrogenic niche anatomy at ureteric bud tips and the role of RET in MAPK activation via ERK. (**B**) Phase contrast micrographs of RET-expressing MDCK cell line as a reductionist model, with and without activation using GDNF and the co-receptor GFRA1. (**C**) Montage of phase and immunofluorescence micrographs and traction force microscopy output (displacement and stress fields) for representative RET-MDCK cells after the indicated treatments on adhesive polyacrylamide substrate. (**D**) Violin plots of traction force distribution for indicated treatments showing means and quartiles (gray lines, *n* = 8 replicate wells per condition; red points are means of replicates). One-way ANOVA, Tukey’s test, \*\*\*\**p* < 0.0001. (**E**) Plots of RET expression and pERK intensity by immunofluorescence upon GDNF/GFRA1 activation vs. untreated control (>1,700 cells combined across *n* = 8 replicate wells). Traction force is represented by point color and by running average plots (window size of 25 cells).

Having validated MATCHY’s TFM integrated with IF, we sought to create a biophysical atlas of the nephron-forming niche from primary mouse embryonic kidney cells (**Fig. 3**). These niches are defined by ‘caps’ of metanephric mesenchyme (nephron progenitor cells) at the tips of the branching ureteric bud (66, 67) (**Fig. 3A**). Nephron progenitors proliferate in the niche and differentiate into early nephron cells that periodically condense by mesenchymal-to-epithelial transition into pre-tubular aggregates (PTAs) (54, 66–69). Such morphological transitions are often triggered by cell rearrangement caused by changes in cell cortical tension and cell-cell adhesion, a process partially explained by models such as the differential adhesion/interfacial tension hypothesis (51–53, 70, 71). Although nephron progenitors express a changing cell adhesion molecule profile as they transition along their differentiation trajectory (10, 22, 24, 25, 69, 72–75), the downstream effect of this on cell biophysical properties has not been quantified. We first verified the molecular basis of nephron MET by analyzing existing mouse kidney scRNA-seq data published by Combes *et al.* (76). Gene set enrichment analysis of a pre-curated list of sets related to cell tension and adhesion revealed significant enrichment for all of them in cells from the PTA and renal vesicle clusters vs. those from the nephron progenitor and ‘primed’ nephron progenitor clusters. Feature plots confirmed differential expression of cadherins *Cdh1, 2, 3, 4, 6*, and *11* over the nephron differentiation trajectory; each being previously identified as relevant to nephron MET and beyond (22, 23, 25, 49, 50).

**Fig. 3:**
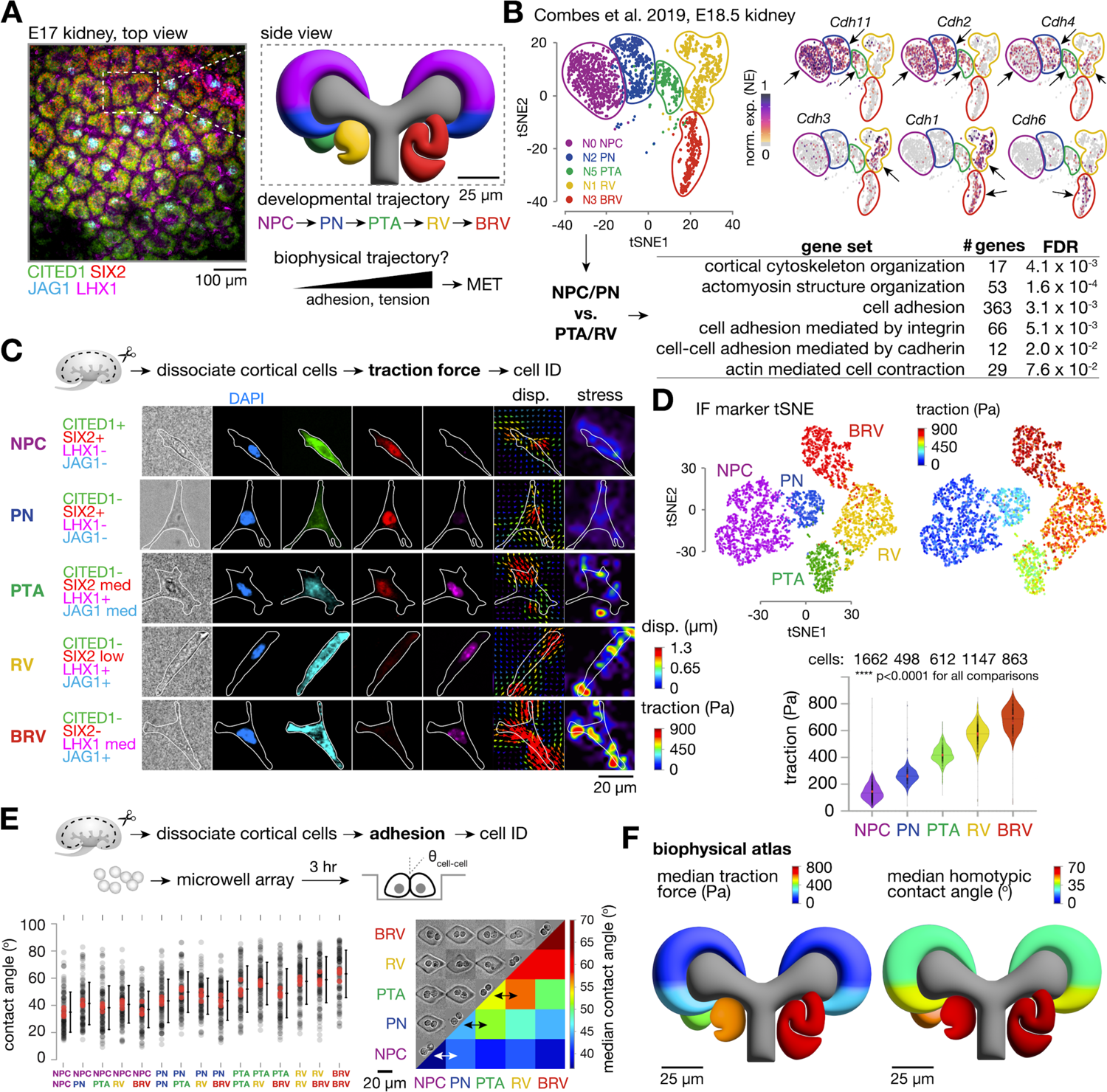
A biophysical atlas of primary mouse nephrogenic niche cells reveals an energetic ratchet accompanying nephron progenitor differentiation. (A) *Left*, Whole-mount immunofluorescence micrograph of E17 kidney cortical surface showing CITED1+ SIX2+ nephron progenitor niche compartments and LHX1, JAG1 differentiation markers. *Right*, schematic of niche anatomy and stages of nephron progenitor differentiation. NPC, nephron progenitor cell; PN, primed nephron progenitor; PTA, pre-tubular aggregate; RV, renal vesicle; BRV, beyond renal vesicle (comma-shaped body, S–shaped body, etc.). (B) *Top*, tSNE plot of cell clusters and feature plots of cadherin expression over the nephron differentiation trajectory from scRNA-seq data published in Combes *et al.* 2019. Arrows indicate clusters having appreciable marker expression. *Bottom*, gene set enrichment analysis results for the listed sets, comparing NPC/PN stages to PTA/RV stages. (C) Montage of phase and immunofluorescence micrographs and traction force microscopy output (displacement, disp., and stress fields) for primary mouse E17 embryonic kidney nephrogenic zone cells representative of each cell type along the differentiation trajectory. (D) *Top*, t-SNE dimensionality reduction plots based on expression of the markers in (C) showing annotation of clusters by cell type (left), and by traction force (right). *Bottom*, Violin plots of traction force by cell type showing means and quartiles (gray lines, *n* = 8 replicate wells; red points are means of replicates). (E) Plot of cell-cell contact angle between the indicated homotypic and heterotypic pairs (mean ± S.D., n > 100 pairs per comparison), and heat-map matrix of median contact angles. Arrows highlight relationship between homotypic contact angle for a given cell type and that for its heterotypic contact with the most closely related cell type along the differentiation trajectory. (F) Biophysical atlas showing median traction and homotypic adhesion data mapped as colors onto the niche schematic. Statistics in (D) and (E) are one-way ANOVA, Tukey’s test, \**p* < 0.05, \*\**p* < 0.01, \*\*\**p* < 0.001.

We next sought to quantify how these molecular-level changes affect cell biophysical properties among differentiating nephron progenitors. Mapping the traction stresses of closely related cell lineages along a differentiation trajectory by traditional means requires them to be separately sorted and assayed, which is challenging or impossible for rare cell types or those having poorly characterized surface marker profiles. Alternatively, lineage-specific reporter mice can be produced to mark cell types of interest by endogenous fluorescence (77). However, this adds significant complexity. We instead recovered ‘nephrogenic zone’ (surface/cortical layer) cells from E17 embryonic mouse kidneys by brief dissociation according to an established protocol (13, 61) and relied on *post hoc* assignment of cell identity after fixation and immunofluorescence for intracellular markers. This cell mixture is enriched for stromal, nephron progenitor, and early nephron lineages. Less than 10% of the recovered cells are mature nephron, ureteric bud, endothelial, or immune cells (13). We performed multiplexed traction force measurements and read out predicted cell identity using thresholds for canonical protein marker expression (**Fig 3C**, nephron progenitor cells, NPC: CITED1+ SIX2+; primed NPC, PN: SIX2+; pre-tubular aggregate, PTA: SIX2+ LHX1+; renal vesicle, RV: SIX2+ LHX1+ JAG1 medium; ‘beyond renal vesicle’, BRV: JAG1+) (78, 79). We then used t-SNE and K-means algorithms, which reduce dimensionality and cluster distinct cell populations (**Fig. 3D**). Overlaying traction data onto these clusters showed a progressive increase along the differentiation trajectory from NPC to BRV. This predicts that MET is associated with an increase in cell contractility, which may be necessary for cell compaction from the niche into PTAs. Indeed, appropriate lumenization of PTAs during their transition to RVs requires non-muscle myosin IIA (*Myh9*) and IIB (*Myh10*) expression, suggesting that cell mechanical tension is likely required for the completion of MET (44). Our data indicate that cells exert higher traction stresses as they progress along the nephron differentiation trajectory.

To complement the model of MET-associated increase in single-cell traction with cell-cell adhesion data, we adapted a microwell-based approach to create highly parallelized arrays of cell doublets (43). We read-out cell adhesion information via contact angle, again inferring cell identities *post hoc* using immunofluorescence (see **Methods**). Similar to our traction data, we measured a monotonically increasing homotypic adhesion along the nephron progenitor differentiation trajectory (**Fig. 3E**). The heterotypic adhesion for a given cell lineage tended to be higher with closely related daughter lineages compared to the homotypic adhesion for that lineage, for example, NPC-PN > NPC-NPC, PN-PTA > PN-PN, PTA-RV > PTA-PTA, and RV-BRV > RV-RV. This structure of increasing heterotypic adhesion to homotypic adhesion may create an ‘energetic ratchet’ that favors physical segregation and transit of cells through the morphological transition to renal vesicles and S-shaped bodies. Intuitively, this ratchet-like process could favor the correct localization and orientation of the forming nephron.

Moreover, cell-cell adhesion appears to reach a maximum at around the RV stage, while traction continues to increase through the BRV stage. This ordering may be necessary to establish sufficient cell-cell adhesion to permit the radical shape change occurring in the nephron upon S-shaped body formation (9, 10, 80). These data indicate a substantial increase in cell-cell adhesion and traction along the differentiation trajectory that may drive structural transition in the developing nephron (**Fig. 3F**).

We next wondered if biophysical modeling of our traction and adhesion data from primary mouse kidney cells would predict self-organization of the niche and early nephrons. Similar models have successfully predicted self-organization of other multicellular structures such as mammary acini (43) and early embryos (40). Previous models for embryonic kidney cell self-organization have partially predicted effects of repulsion/attraction within and between the cap mesenchyme and the ureteric bud on bud/niche organization, but did not quantitatively measure cell biophysical properties or attempt to explain self-organization of nephron lineage cells during early nephrogenesis (22, 81, 82). To address this question, we adapted a cellular Potts (agent-based) model for early embryo self-organization (40); drawing cell-cell adhesion and cortical tension parameters from our quantitative biophysical atlas distributions (**Fig. 3D-F**, see **Methods**). We hypothesized that the measured biophysical parameters from our experiments on dissociated primary nephrogenic zone cells would be sufficient to explain nephron condensation, location in the niche, and proximal-distal polarization (i.e. spatial ordering of cells along the differentiation trajectory during segment patterning, **Fig. 4A**).

**Fig. 4:**
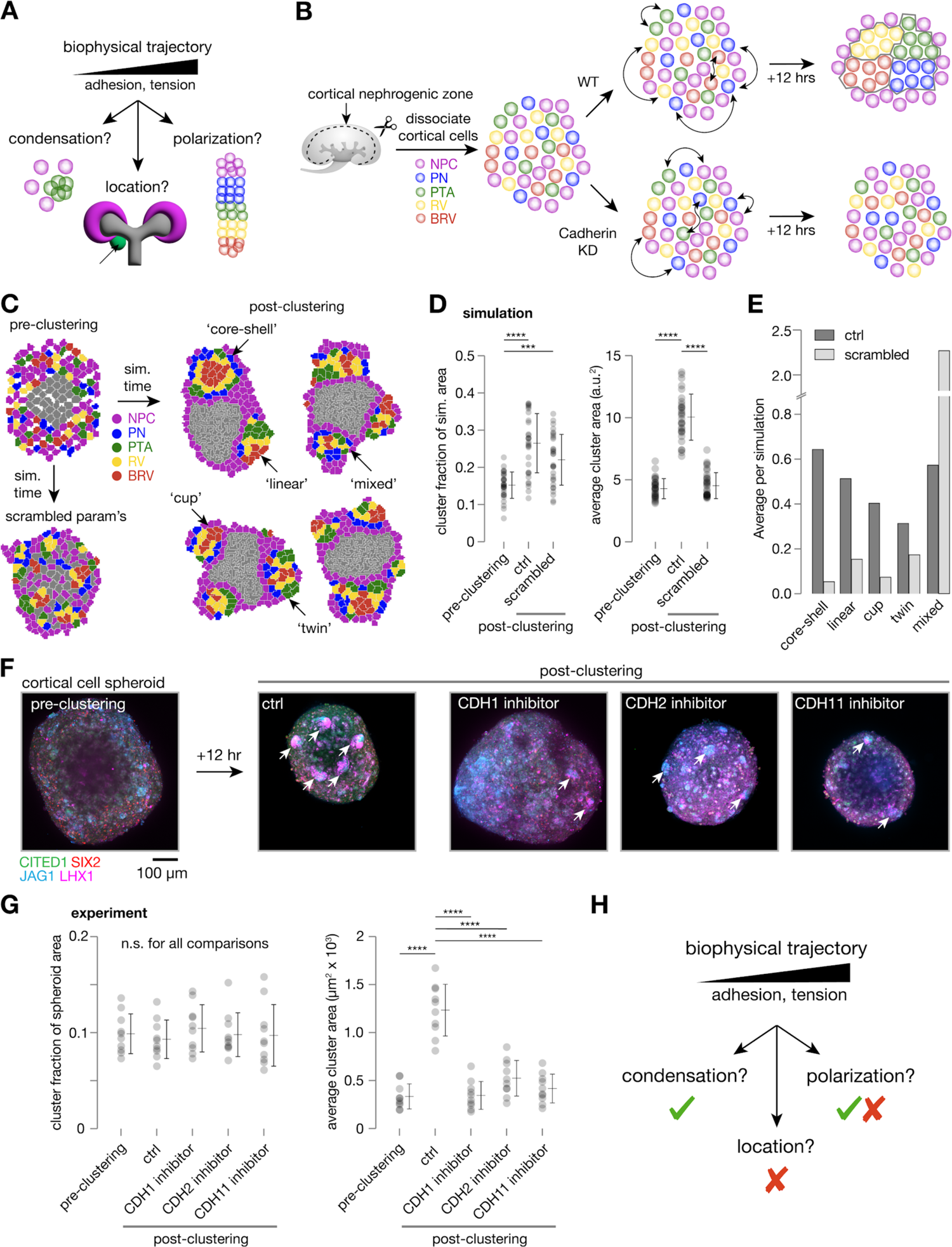
An energetic ratchet in cell contractility and adhesion properties is sufficient to explain early nephron condensation and partially sufficient to explain subsequent polarization. (A) Schematic of nephron formation properties potentially affected by an energetic ratchet in cell-cell adhesion and tension during nephron progenitor commitment. (B) Schematic of hypothesis for dissociated mouse embryonic kidney cell self-organization driven by cell biophysical properties. (C) Agent-based simulations of nephrogenic niche self-organization. *Top left*, Example simulation initial condition with ureteric bud cells clustered together and other nephron lineage cells randomly arranged around it. *Bottom left*, example ‘scrambled’ simulation with biophysical properties of cells randomly chosen from all traction and adhesion distributions. *Right*, example control (ctrl) simulations with biophysical properties of cells randomly chosen from the traction and adhesion distributions corresponding to the correct cell lineage. Condensed structures are qualitatively annotated according to the indicated morphology types: ‘core-shell’ (annular distribution of cell types along the differentiation trajectory), ‘linear’ (linear distribution), ‘mixed’ (no apparent order), ‘cup’ (combining features of core-shell and linear), or ‘twin’ (clusters with two linear axes fused with each other). (D) Plots of fraction of simulated nephrogenic niche occupied by clusters, and average individual cluster area per spheroid (points are simulation instances, mean ± S.D., n > 30 simulations per condition). (E) Plot of representation of qualitative sorting states in the model. (F) Immunofluorescence micrographs of primary E17 nephrogenic zone cell spheroids immediately after aggregation (*left*), and after 12 hr incubation in control media (ctrl) or in the presence of blocking antibodies for the indicated cadherins (*right*). Arrows indicate condensed cell clusters. (G) Plots of fraction of spheroid area occupied by clusters, and average individual cluster area per spheroid (points are spheroids, mean ± S.D., n > 10 spheroids per condition). (H) Schematic summary of biophysical prediction of nephron formation properties. Statistics in (D) and (G) are one-way ANOVA, Tukey’s test, \*\*\**p* < 0.001, \*\*\*\**p* < 0.0001.

We paired the model with a self-organization assay consisting of spheroid cultures of the same primary cells (**Fig. 4B**). Previous work has noted intimate interactions between cap mesenchyme (nephron progenitor) cells and the ureteric bud mediated by adhesion molecules such as ITGA8 (76, 83). However, we did not recover ureteric bud cells and measure their traction or adhesion properties. Instead we assumed that their homotypic adhesion and heterotypic adhesion with nephron progenitors were arbitrarily high (since ureteric bud cells form core-shell structures with nephron progenitors in re-aggregation assays (22, 46)). We also assumed that the heterotypic adhesion between ureteric bud cells and more differentiated nephron lineages was negligible (since these never mix *in vivo*). Using cell-cell adhesion and traction force parameters drawn from our biophysical atlas in the simulation led to spontaneous clustering and cell organization given initial seeding conditions in agent position and composition similar to that in the mouse nephrogenic niche (**Fig. 4C,D**, **Movie S1**). Differentiated agents that were initially randomly distributed condensed and formed several qualitative morphologies roughly reminiscent of early polarization of nephrons at the renal vesicle and S-shaped body stages (**Fig. 4C,E**). Both clustering and partial polarization phenotypes were significantly less frequent when parameters were ‘scrambled’ by randomly selecting values from the measured cell type distributions. Mimicking this, 12 hr culture of our primary cell spheroids was sufficient to observe clustering of LHX1+ JAG1+ early nephron cells, which was significantly ablated in the presence of blocking antibodies for CDH1 (expressed in RV, BRV), CDH2 (NPC, PN, PTA), or CDH11 (NPC, PN, PTA) (22, 23, 49, 50, 76, 84) (**Fig 3B**, **Fig. 4F,G**). These data conflict with findings that genetic knockout of individual cadherins does not significantly undermine mouse nephron MET in CHIR-induced primary nephron progenitor cell cultures (specifically *Cdh2, 3, 4,* or *11*) or mouse models (*Cdh4*^-/-^, *Cdh6*^-/-^) (24, 49, 50). However, knockout of *Cdh6* (and potentially of *Cdh4*) does delay nephron epithelialization (24, 50). Furthermore, kidney explant culture magnifies the loss of MET in *Cdh4*^-/-^ kidneys and when a blocking antibody for CDH6 is used in wild-type kidneys (23, 50). These data suggest that our results may arise from the relatively brief self-organization period assayed here and/or other properties of our *in vitro* culture approach (e.g. condensation of already differentiated cells rather than of nephron progenitors undergoing induction). Overall the primary cell and modeling data indicate that early nephron condensation and some of the spatial structure of the mouse nephrogenic niche can be explained on the basis of nephron lineage cell biophysical properties alone. Second, the energetic ratchet likely locally reinforces or provides an error-correcting function to other contributions to especially nephron polarization, which is thought to require additional factors provided by the ureteric bud and surrounding stromal cells (19, 78, 85–87).

## Discussion

Gene expression and signaling pathways operate through single-cell and supracellular biophysical properties to determine tissue structure (88). A quantitative understanding of biophysical changes along differentiation trajectories is therefore necessary to guide tissue organization in engineered, developing, and diseased tissues alike. Cell-cell heterogeneity necessitates high-throughput measurements to capture the full distribution of biophysical changes within and between multiple cell states. Here we address these needs in MATCHY via microfabrication, serial integration of biophysical and molecular characterization assays, and machine learning automation. We achieve quantitation of adhesion and traction of >10,000 cells across >8 independent experimental conditions in the same experiment, all within 12 hr of tissue dissociation. The MATCHY approach is adaptable to any tissue type amenable to single-cell dissociation. We chose kidney development as a case study due to the complexity of cell dynamics, decision-making, and self-organization within its nephrogenic niches. We first validated MATCHY performance for GDNF-RET tyrosine kinase signaling in a cell line model of its effects on ureteric bud branching morphogenesis, revealing a correlation between MAPK signaling activation and downstream cell traction that scales with RET receptor expression. We then explored the multiplexing capability of MATCHY for primary cell suspensions prepared from mouse embryonic kidneys, finding monotonic increases in both cell traction and adhesion along the nephron differentiation trajectory that complement previous characterization of an expression-level adhesive switch program associated with MET. The adhesion data reveal an ‘energetic ratchet’ where the heterotypic adhesion of a given cell type with its most closely related differentiation state is higher than its homotypic adhesion. This would tend to spatially recruit differentiating cells into progressively more mature tissue compartments, potentially explaining physical segregation of newly formed nephrons from the niche. To explore this, we performed agent-based modeling, which predicted condensation of early nephrons in the nephron progenitor niche using parameters sampled from cell experiments. The model also partially predicted spatial sorting of differentiation states in an analogous fashion to that occurring during polarization of the nephron *in vivo*. This highlights the likely need for additional ureteric bud and/or stromal derived factors to increase the likelihood of correct linear organization of nephrons and in the correct orientation along the proximal-distal axis. Together these data provide a biophysical atlas of nephrogenesis, an important roadmap for tissue engineering efforts to reconstitute nephrogenesis in iPSC-derived kidney organoids for regenerative medicine applications.

Our contributions here leave several areas for future study and consideration. Firstly, we did not consider traction or adhesion properties of ureteric bud or stromal cells. These compartments form important niche boundary interfaces that likely contribute to nephron progenitor sorting dynamics. For example, the ureteric bud makes adhesive interactions with nephron progenitors through a range of cell-cell and cell-matrix ligand-receptor pairs, notably ITGA8-nephronectin, which is required for proper niche organization (45, 76, 83). Similarly, the underlying cause of new nephron positioning at the curved ‘armpits’ of branching ureteric bud tips is an active area of interest (89, 90). Our simulations show some intriguing curvature of the ureteric bud local to sites of nephron condensation, suggesting that this may be more of a ‘chicken and egg’ problem than previously recognized. The renal stroma, which forms ‘ribbons’ that divide niches and surround newly forming nephrons, was recently shown to have significant spatial heterogeneity local to the niche and along the cortico-medullary axis (91, 92), and basal adhesion of cells to surrounding stroma may contribute to sorting outcomes (43). The contribution of ureteric bud and stromal cells could be readily integrated in future work. Second, removing cells from their native tissue environment risks distorting readout of true *in vivo* biophysical properties by altering surface adhesion receptor integrity and limiting measurements to those possible through cell interactions with an appropriate traction force substrate or single partner cell type in adhesion measurements. For many questions, these caveats are likely to be a reasonable tradeoff against the volume of information that can be gathered by MATCHY compared to the few *in vivo* measurement tools that exist (37, 93–99).

In summary, MATCHY contributes a high-throughput and user-friendly extension to existing cell biophysical characterization techniques. MATCHY provides a powerful benchmarking tool for mechanobiology studies and emerging tissue engineering strategies. These seek to synthetically control and leverage self-organization principles to build complex tissue ‘seeds’ with further developmental potential (27, 40, 100). In the kidney engineering space, for example, such tight feedback between multicellular design, biophysical measurement, structure prediction, and *in vitro* reconstitution carries a promising potential to control nephron formation, polarization, and connectivity. Kidney tissues assembled by leveraging biophysical principles have enormous potential to contribute a third arm to kidney replacement strategies beyond transplantation and dialysis in the future.

## Acknowledgements

We are grateful to Gregory Dressler (University of Michigan) for the generous gift of MDCK-RET9 cells. We thank John Viola and Samuel Grindel for sharing their expertise on experiment protocols and spheroid segmentation. This work was carried out in part at the Singh Center for Nanotechnology at the University of Pennsylvania supported by the NSF National Nanotechnology Coordinated Infrastructure Program (NNCI-2025608). This research was partially supported by the NSF through the University of Pennsylvania Materials Research Science and Engineering Center (MRSEC, DMR-2309043). This work was supported by an NSF GRFP award (JL), NIH F32 DK126385 (LSP), NIH NIGMS MIRA R35GM133380 (AJH), NIH NIDDK R01DK132296 (AJH), and NSF CAREER award 2047271 (AJH).

## Supplementary Information

### Supplementary Figures

**Fig. S1:**
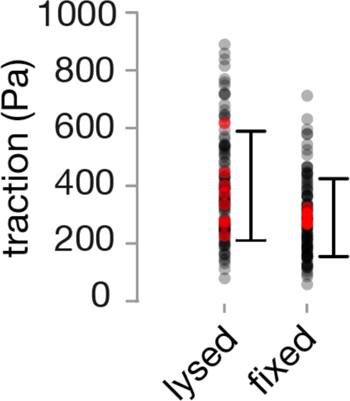
Cell fixation during TFM recovers the majority of traction magnitude relative to lysis. Plot of cell traction for iPSCs measured relative to a relaxed TFM gel state generated by cell lysis or fixation (>100 cells across *n* = 8 replicate wells per condition; red points are means of replicates).

### Supplementary Movies

**Movie S1: Agent-based simulations of nephron lineage self-organization.** Movie of simulation dynamics for example ‘scrambled’ and control (‘ctrl’) instances over 10^7^ simulation steps.

## Methods

### Human iPSC cell culture, distal nephron and ureteric bud epithelium derivation

Distal nephron cells (DN) and ureteric bud epithelium (UB) were generated from GATA3^YFP^ and GATA3^mCherry^ transgenic reporter iPSC lines respectively (*GATA3-T2A-optoWnt-T2A-YFP*, *GATA3-T2A-mCherry* Murdoch Children’s Research Institute / Kidney Translational Research Center, Washington University Nephrology) according to published protocols (101, 102). Briefly, iPSCs were maintained in standard tissue-culture treated 6-well plates in stem cell maintenance medium plus supplement (mTeSR+ kit, StemCell Technologies 100-0276). Cells were passaged using Accutase (StemCell Technologies, 07920) and plated at a density of 100k cells/well. Cells were maintained at a confluency below ∼70%.

A day prior to DN production, cells were checked for off-target differentiation before plating at 200k cells/well. Full media was used to culture the cells for 24 hr before being replaced with TeSR-E6 Medium plus supplements (TeSR-E6+) (STEMCELL Technologeis 05946) and 7μM CHIR 99021 (Torcis, 4423) for 4 days. On day 5, the media was changed to TeSR-E6+, 1 μg/ml heparin (Sigma-Aldrich H4784), 600 ng/ml FGF2 (PeproTech, 100-18B), and 1 μM CHIR for 3 days. On day 8, cells were passed at a 1:1 ratio into a ultra-low attachment 6-well plate (Corning, CLS3471) placed on an orbital shaker at 60 rpm in a standard cell culture incubator at 37°C and 5% CO_2_. The same media plus 0.1% PVA (Sigma, 182508), 0.1% MC (Sigma, M0512), 10 μM Rho kinase inhibitor (RI, Y-27632) (Tocris, 1254) was used. After 24 hr, the media minus RI was used for further culture for another 4 days before cells were FACS purified.

For UB derivation, one day before differentiation cells were plated at 50k/well on standard tissue-culture treated 6-well plate. After 24 hr, TeSR-E6+ and 7μM CHIR were used for 4 days and 600 ng/ml FGF2 plus 1μg/ml heparin for 3 days before cells were dissociated and suspended at 200k cells/μl. Cell ‘pucks’ were made by dispensing 3 μl of the suspension on a 6-well transwell plate with 1 ml TeSR-E6+ with 7 μM CHIR added to the lower compartment. After 1 hr of incubation, the media was changed to 600 ng/ml FGF2 plus 1 μg/ml heparin for 5 days and refreshed every second day. The media was then switched to TeSR-E6+ with 0.1 μM TTNPB (Tocris, 0761) for another 9 days, during which 1 ml media was added to the lower compartment and refreshed every second day.

GATA3-YFP+ DN and GATA3-mCherry+ UB were dissociated and FACS purified at the end of differentiations. Cell suspensions were passed through a 40 μm cell-strainer, aggregated, and resuspended at 1ml cells/ml in PBS with 2% FBS. Positive cells were sorted with a 100 μm nozzle at ∼600 events/s flow rate. Cells were collected at the end of the purification and resuspended in TeSR-E6+ before use in experiments.

### MDCK cell culture, cell suspension preparation, and perturbations

Madin-Darby canine kidney (MDCK) epithelial cells were maintained in minimum essential medium (MEM, Earle’s salts and L-glutamine, #MT10-010-CM, Corning) supplemented with 10% FBS and 100 U ml^-1^ penicillin-streptomycin, and were passaged using 0.25% trypsin-EDTA (#25200056, Corning). MDCK-hRET9 and MDCK-hRET9^KM^ cell lines (55) were routinely cultured in a selection medium containing 100 µg ml^-1^ neomycin (G418, 50 µg ml^-1^ stock, Penn Cell Center Stockroom) to remove non-expressing cells. Live imaging medium for cell lines consisted of phenol red-free DMEM (4.5 g l^-1^ glucose, L-glutamine, and 25 mM HEPES, #21063-029, Invitrogen) supplemented with 10% FBS, 1 mM sodium pyruvate (100 mM stock, #11360070, Invitrogen), and 100 U ml^-1^ penicillin-streptomycin. All cell lines were subcultured in polystyrene flasks that were maintained in a humidified incubator at 37°C and 5% CO_2_.

Prior to cell seeding for contractility characterization, MDCK cell cultures were washed with DPBS for 1 min before treatment with 0.25% trypsin-EDTA for 10 min. Full medium was used to terminate the digestion and cell pellets were aggregated with microcentrifugation at 300 g for 3 min. Cell suspensions were reconstituted at 400,000 cells/ml with 0.5 ml being seeded per well in an 8-well chambered slide. To stimulate pERK overexpression, 100 ng/ml of soluble, recombinant human GFRA1 (R&D Systems, AF714-SP) along with 50 ng/ml recombinant human GDNF (R&D Systems, 212-GD-010) were administered for 12 hr. ROCK inhibitor (STEMCELL Technologies, 72304) and blebbistatin (Tocris, 1852) treatments were carried out at 10 µM and 50 µM, respectively.

### Animal experiments

Mouse protocols followed NIH guidelines and were approved by the Institutional Animal Care and Use Committee of the University of Pennsylvania. E17 embryos were collected from timed pregnant CD-1 mice (Charles River) and stages confirmed by limb anatomy as previously described (103). Embryonic kidneys were dissected in chilled Dulbecco’s phosphate-buffered saline (DPBS, MT21-31-CV, Corning) (104).

### Kidney immunofluorescence imaging

Immunofluorescence staining and imaging was performed as previously described (105), using protocols adapted from Combes *et al.* and O’Brien *et al.* (83, 106). Briefly, dissected kidneys were fixed in ice cold 4% paraformaldehyde in DPBS for 20 min, washed three times for 5 min per wash in ice cold DPBS, blocked for 2 hr at room temperature in PBSTX (DPBS + 0.1% Triton X-100) containing 5% donkey serum (D9663, Sigma), incubated in primary and then secondary antibodies in blocking buffer for at least 48 hr at 4°C, alternating with 3 washes in PBSTX totaling 12-24 hr. The minimum duration of primary and secondary incubations and washes depended on the age of the kidney, as previously described (106). Primary antibodies and dilutions included rabbit anti-Six2 (1:600, 11562-1-AP, Proteintech, RRID: AB_2189084), goat anti-E-cadherin (1:200, AF748, R&D systems, RRID: AB_355568), rat anti-PDGFRA (1:200, 14-1401-82, Invitrogen, RRID: AB_46749), goat anti-ITGA8 (1:20, AF4076, R&D systems, RRID: AB_2296280), rabbit anti-LEF1 (1:200, 2230, Cell Signaling Technologies, RRID: AB_823558), rat anti-E-cadherin (1:100, ab11512, abcam, RRID: AB_298118), mouse anti-E-cadherin (1:200, clone 34, 610404, BD Biosciences, RRID: AB_397787), rabbit anti-MYL12A (phospho S19) (1:100, ab2480, abcam, RRID: AB_303094), goat anti-jagged 1 (1:150, AF599, R&D Systems, RRID: AB_2128257), rabbit anti-RET (1:200, 3223, Cell Signaling Technologies, RRID: AB_2238465). Secondary antibodies (all raised in donkey) were used at 1:300 dilution and include anti-rabbit AlexaFluor 405 (ThermoFisher, A48258, RRID: AB_2890547), anti-rat AlexaFluor 555 (ThermoFisher, A48270, RRID: AB_2896336), anti-goat AlexaFluor 488 (A11055, ThermoFisher, RRID: AB_2534102), anti-rat AlexaFluor Plus 405 (A48268, ThermoFisher, RRID: AB_2890549), anti-rabbit AlexaFluor 555 (A31570, ThermoFisher, RRID: AB_2536180), anti-goat AlexaFluor 647 (A32849, ThermoFisher, RRID: AB_2762840), anti-mouse AlexaFluor 405 (A48257, ThermoFisher, RRID: AB_2884884), anti-rabbit AlexaFluor 488 (A21206, ThermoFisher, RRID: AB_2535792). In some experiments, samples were counterstained in 300 nM DAPI (4’,6-diamidino-2-phenylindole; D1306, ThermoFisher) diluted in blocking buffer for 2 hr at room temperature, followed by 3 washes in PBS.

Kidneys were imaged in wells created with a 2 mm diameter biopsy punch in a ∼5 mm-thick layer of 15:1 (base:crosslinker) polydimethylsiloxane (PDMS) elastomer (Sylgard 184, 2065622, Ellsworth Adhesives) set in 35 mm coverslip-bottom dishes (FD35-100, World Precision Instruments). Imaging was performed using a Nikon Ti2-E microscope equipped with a CSU-W1 spinning disk (Yokogawa), a white light LED, laser illumination (100 mW 405, 488, and 561 nm lasers and a 75 mW 640 nm laser), a Prime 95B back-illuminated sCMOS camera (Photometrics), motorized stage, 4x/0.2 NA, 10x/0.25 NA and 20x/0.5 NA lenses (Nikon), and a stagetop environmental enclosure (OkoLabs).

### Primary mouse embryonic kidney dissociation

Primary embryonic mouse nephrogenic whole niche cell suspensions were made by adapting previously described protocols (13, 107, 108). Briefly, a digest solution consisting of 0.5% pancreatin (Sigma-Aldrich, P7545) and 0.25% Collagenase A (Sigma-Aldrich, C0130) in DPBS was prepared. Pancreatin was first added to DPBS and allowed to solubilize on a rotator at 4°C overnight. Collagenase was added to the digest solution and the mixture was solubilized on a rotator at room temperature for 3-5 hr on the day of the experiment. Batches of 8 dissected E17 embryonic kidneys were incubated in 1.5 ml digest solution for 15 min at 37°C. The digest reaction was then stopped by adding 75 µL of fetal bovine serum. 1.4 ml of the digest solution containing dissociated cells was then transferred to a fresh microcentrifuge tube containing 2 µL of DNase (Fisher Scientific, AM2238) and incubated for 10 min at 37°C. The cell solution was centrifuged and resuspended 2 times in 1 ml Hank’s Balanced Salt Solution (HBSS) + 5% fetal bovine serum. Cells were then resuspended in media and passed twice through 40 µm cell strainers (CLS431750, Corning) by pipetting.

### MATCHY traction force microscopy

A 100 mm-diameter mechanical-grade silicon wafer (University Wafer) was fabricated with shims for precise gel spacing as previously described (109). Briefly, a ‘shim’ photomask reflecting the geometry of a standard 1’’ x 3’’ microscope slide (12-559-A3, ThermoFisher) was printed at 20,000 d.p.i (CAD/Art Services) to cast parallel rails on the wafer to allow 30 µm-thick gel manufacturing later. The wafer was first baked on a hot plate at 200°C for 10 min and then centered on a vacuum spin coater (SCK-300P, Instras Scientific). Microchem SU-8 2025 photoresist (Fisher Scientific) was dispensed on the wafer at 4 ml and spin-coated at 500 rpm for 12 s followed by 3,000 rpm for 45 s to create a uniform 30 µm layer of photoresist. The assembly was then baked on a hot plate at 95°C for 8 min. The photomask was placed on the treated wafer and exposed to 365 nm ultraviolet light at 175 mJ/cm^2^ (M365LP1, Thorlabs). It was further baked at 95°C for 8 min and developed by submerging in SU-8 developer (Microchem SU-8 developer, Fisher Scientific) for at least 15 min. The wafer was rinsed with acetone and isopropanol and left to air dry. Vapor deposition was used to render the wafer hydrophobic with 1 ml of dichlorodimethysilane (DCDMS, Sigma).

Standard microscope slides were functionalized for polyacrylamide (PA) gel covalent attachment by etching in 1N sodium hydroxide (NaOH, S8045, Sigma) for 15 min and silanizing with 3-(trimethoxysilyl)propyl methacrylate (440159, Sigma), acetic acid (EMD Millipore), and deionized (DI) water at 2:3:5 ratio by sandwiching 150 µL solution between two slides. The ‘slide sandwiches’ were incubated for at least 30 min at room temperature and rinsed with 100% methanol and DI water.

The silanized side of the treated slide was placed face down on top of the shims on the wafer. PA gel precursor solution was prepared by pipetting 645.5 µL DI water, 187.5 µL acrylamide solution (40% w/v Acrylamide Solution, 1610140, Bio-Rad), 100 µl 10x DPBS (14200075, Thermo Fisher Scientific), 35 µL N,N-methylenebisacrylamide crosslinker (2% w/v bis-acrylamide Solution, 1610142, Bio-Rad), 20 µL of 1:100 diluted fluorescent beads in DI water (FluoSpheres carboxylate-modified 0.2 µm, 660/680, F8807, Thermo Fisher Scientific), 6 µL 10% v/v tetramethylethylenediamine (TEMED) (Sigma) and 6 µL 10% w/v ammonium persulfate (Sigma) in DI water. PA gels formed from this precursor have a Young’s modulus of 6 kPa (58). The precursor was degassed under vacuum in an ultrasonic cleaning bath (15-337-411, Thermo Fisher Scientific), and 120 µL was dispensed in the gap between the slide and wafer and allowed to cure for 30 min protected from light exposure.

To derivatize uncoupled acrylamide chains on the gel surface with NHS ester for adhesive protein functionalization, 0.01% (w/v) N,N-methylenebisacrylamide in 50 mM (pH 6.0) 4-(2-hydroxyethyl)-1-piperazineethanesulfonic acid (HEPES, 40820004, bioWORLD) buffer was mixed with 0.09% (w/v) lithium phenyl-2,4,6-trimethylbenzoylphosphinate (LAP photoinitiator, 900889, Sigma) and 0.1% (w/v) acrylic acid-*N*-hydroxysuccinimide ester (N2 acrylic acid, A8060, Sigma), as previously reported (110). Free-radical photochemistry was carried out by pipetting 200 µl of the mixture on top of the PA gel and initiated under UV-light (I365 nm = 15 mW/cm^2^) for 30 min. The NHS-coated PA gel was washed in ice-cold DI water and 100 µM sodium chloride (NaCl, S6191, Sigma) and assembled into a clip-on 8-well chambered slide with a silicone gasket (CCS-8, MatTek). The wells were incubated in 100 mg/ml Matrigel (356234, Corning) overnight at 4°C to covalently couple the N-terminus of the ECM proteins before washing slides four times with PBS at room temperature. After the last wash, primary cell culture media was used to incubate the wells before cell seeding.

Cell suspensions from primary tissue or cell culture plates (see **Primary mouse embryonic kidney dissociation; MDCK cell culture, cell suspension preparation, and perturbations**) were seeded at 20,000 cells per well and cultured for 12 hr to allow attachment and traction equilibration. Brightfield and Cy5 confocal images of cell and bead positions were then captured in the full-traction state. An automated *xy* point assignment program was populated to record user-defined multipoints within each well, and autofocus was used to automate collection of *z*-stacks.

To read out the traction forces, cells were fixed in 4% paraformaldehyde (16% PFA, 15710, Electron Microscopy Sciences) in 1x DPBS for 20 min and washed with DPBS four times at 5 min each. The wells were loaded on the confocal microscope and global registration performed by manually fine-tuning the four corners of the slide to align the same positions previously recorded in the full-traction state. After global translational and rotational displacements were corrected, the same *xy* points were imaged. Autofocus was used to capture *z*-stacks of the beads identical to those of the full-traction state.

For downstream immunofluorescence, cells were permeabilized in 0.5% v/v Triton X-100 for 5 min in 1x DPBS and blocked with 1% BSA in IF wash buffer (1% w/v BSA, 2% v/v Triton X-100; 0.41% v/v Tween-20, EMD Millipore, 1x DPBS) for 1 hr.

The wells were then incubated with fluorophore-conjugated primary antibodies: rabbit anti-Six2 (11562-1-AP, Proteintech, RRID: AB_2189084) labeled with Alexa Fluorphore 555 (Novus, 333-0010), mouse anti-Lhx1 (CF504527, OriGene, RRID: AB_2724601) labeled with Alexa Fluorphore 647 (Novus, 336-0010), and rabbit Cited1-488 (Proteintech, CL48826999100UL, RRID: AB_2919211) as the first set and rabbit anti-SIX2-555, mouse anti-LHX1-647, and goat anti-JAG1 (AF599, R&D Systems, RRID: AB_2128257) labeled with Alexa Fluophore 488 (Novus, 332-0005). All primary antibodies were firstly cleaned up using an antibody purification kit (Novus, 861-0010) and then conjugated with fluorophores according to user manuals. All conjugated antibodies were used at 1:50 dilution overnight at 4°C. Cells were then washed with IF buffer 3 x 1 hr each while protected from light, and nuclei were counterstained with 150 nM DAPI for 15 min. F-actin was stained with 1X Phalloidin-555 (Abcam, ab176756) in PBS + 1% BSA for 1 hr. Finally, wells were washed with IF buffer and PBS for 1 hr each before confocal immunofluorescence imaging of antibody markers using 20 µm *z*-stacks above the gel surface after a similar *xy* registration procedure.

Image analysis was performed using automated macros in ImageJ/Fiji (111). “Find Focused Slides” were used to select the corresponding *z*-frames between the full-traction and relaxed states, and z-stack maximum projections were concatenated to run registration with the “Linear Stack Alignment with SIFT” plugin (112). Displacement and traction fields were calculated with the PIV and FTTC plugins respectively (60). Automated cell segmentation was carried out with the StarDist plugin (113) and only single cells were analyzed. Specifically, the DAPI channel was used to segment cell nuclei, and cells were excluded if their diameters were smaller than 5 µm or larger than 20 µm. If the Euclidean distance between two nuclei was smaller than twice the sum of their major axes, both cells were excluded. The filtered ROIs were then looped through to extract the mechanical and biomolecular information of the cells. A 60 x 60 µm window centered on each cell was drawn to calculate the average traction force and pixel intensity, counting the top 20th and 10th percentiles of the pixels, respectively. The cell ID number along with its traction and IF data was stored before further processing. For cell identification, 50 positive cells of each nephron subpopulation were manually identified based on the known canonical gene expression patterns, and their biomarker expression levels were individually recorded in terms of fluorescence intensity. Fluorescence cutoffs were determined by analyzing the intensity ranges of different cell types and setting thresholds that distinguished unique populations. A lower and upper cutoff values of each marker gene were determined for different cell types to be referred to for downstream analysis. Dimensional reduction of biomarker levels were performed using t-distributed stochastic neighbor embedding (tSNE) in MATLAB to derive clusters. Each nephron lineage was assigned a unique identity by cross-checking with the previously determined threshold values and referring to canonical gene expression patterns from literature.

### MATCHY cell-cell contact angle/adhesion measurement

A contact angle assay was adapted from Cerchiari *et al.* to allow for downstream multiplexing capability at high-throughput (43). To produce microwells hosting cell doublets, silicon wafer fabrication and PA gel casting were performed similarly as described in **MATCHY traction force microscopy**. While reflecting the geometry of a standard 1” x 3” microscope slide, the ‘shim’ photomask additionally contained 20 x 40 µm features throughout the printed area at 20,000 d.p.i (CAD/Art Services) to cast parallel rails as well as microwells. The remaining photolithography and PA gel casting approaches remained the same to fabricate 30 µm-thick microwells. Cell suspensions from primary tissue (see **Primary mouse embryonic kidney dissociation**) were seeded at 10^6^ cells per well in 500 µl final culture media on an 8-well chambered slide with microwells. The cells were settled for 5 min by gravity into the microwells, while being monitored intermittently during the process. Once the majority of the microwells were occupied, the microwells were gently washed with DPBS three times and with media twice. Each well was supplemented with 500 µl culture media and incubated for 3 hr at 37°C before imaging. Imaging was performed using a 20x objective; rapidly capturing doublets by loading pre-calculated *xy* coordinates given known nominal locations of microwells. Imaging was completed within an additional 2 hr (i.e. within 5 hr from doublet seeding).

Cells were immediately fixed and stained after imaging. The same *xy* coordinates were used to capture the doublets again for identification given marker expressions by thresholding (see **Traction force microscopy**). To measure contact angle, the angle tool in ImageJ was used to measure the average of 4 measurements of contact angle θ_cell-cell_ per cell doublet (see schematic in Fig. 3E).

### Primary embryonic mouse kidney whole nephrogenic niche spheroids

Dissociated E17 cells were plated at 50,000 cells per well in 96-well ultra low attachment round-bottom plates (Corning 7007) in Advanced RPMI 1640 Medium and Glutamax (see **Primary mouse embryonic kidney dissociation; MDCK cell culture, cell suspension preparation, and perturbations**). For cadherin blocking conditions, E-cadherin (Cdh1) antibody (1:10, 13-1800 Thermo Fisher, RRID: AB_2533004), N-cadherin (Cdh2) antibody (1:20, 13-2100, Thermo Fisher, RRID: AB_86579), and Cadherin-11 (Cdh11) antibody (1:10, AF1790, RRID: AB_2076971) were administered. Cells were aggregated by plate centrifugation at 300 g for 30 sec and transferred to a standard tissue culture incubator. The control spheroids were harvested 2 hr after aggregation, fixed, and immunofluorescence stained for segmentation (see **Kidney immunofluorescence imaging**). The experimental spheroids as well as those with perturbations were incubated for 12 hr before fixation and staining. Cluster size quantification was performed in ImageJ using the *Threshold* and *Analyze Particles* functions.

### Cellular Potts Model

A Cellular Potts Model (CPM) implemented in Python was adapted from Bao *et al.* to predict cell sorting given our biophysical measurements of cell contractility and adhesion (40). In brief, we parameterized contractility and adhesion strengths using our direct TFM and contact angle measurements and sampled these distributions to build the contractility term (*ƙ*) and the adhesion matrix (*J*). For the CPM simulation, a randomly selected contractility energy (*ƙ*) of the correct cell type and sampled *J* matrix that contained adhesion energies between all appropriate pairs of cell types were used to generate the sorting configuration over time. For the control CPM simulation, the contractility term was randomly sampled among cell types and the *J* matrix was built by sampling from scrambled adhesion energy between cell pairs, thereby randomly assigning the energy terms to different populations. The energy function in **Eq. 1** from Bao *et al.* (40) was minimized via a stochastic process that takes into account both differential affinity and physical properties of the cells.

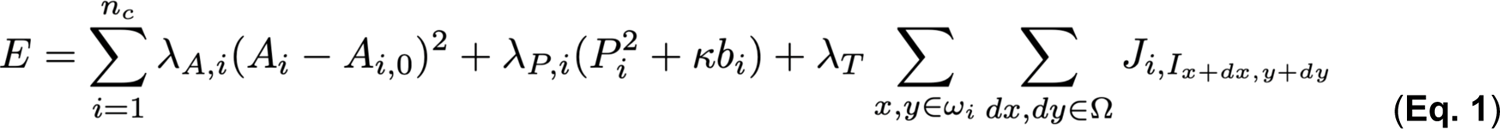

This equation captures cell area penalty (left term), contractility/cortical tension (middle term), and homotypic/heterotypic adhesion (right term). Terms are defined in Bao *et al.* (40). Briefly:

λ_A,i_: Bulk modulus of area deformation of cell *i* from an optimum

A*_i_*_,0_. λ_P,i_: Circumferential elastic modulus of the perimeter.

P*_i_*^2^: Contractility term.

*κb_i_*: Interfacial tension between cells and media (where *b_i_* is the number of Moore neighbors of cell *i* that are medium).

ωi: Lattice points x,y that the cell occupies

Ω: Moore neighborhood such that I*_x+dx,y+dy_* is the cell ID of a lattice point neighboring a point within the cell.

J_i,I*x+dx,y+dy*_: Interaction strength between cell *i* and the neighborhood cell.

λ_T_: Scale factor for adhesion terms.

Relevant cell types were initialized at a similar *in vivo* ratio to that in the mouse nephrogenic niche with 60 NPC, 15 PN, 15 PTA, 15 RV, 15 BRV, and 30 UE cells per simulation. The modeling outcomes were quantified by measuring clustering sizes of contiguous cell connections of more than 2 cells of the same cell types or scored as being one of 5 sorted configurations by qualitatively assigning each cluster to ‘core-shell’ (annular distribution of cell types along the differentiation trajectory), ‘linear’ (linear distribution), ‘mixed’ (no apparent order), ‘cup’ (combining features of core-shell and linear), or ‘twin’ (clusters with two linear axes fused with each other) categories.

### Single-cell RNA sequencing data analysis

Gene expression matrices generated from scRNA-seq of dissociated E18.5 mouse embryonic kidneys in Combes *et al.* were used for analysis (76). The expression matrices of three kidney samples were accessed from the Gene Expression Omnibus (GEO, NCBI) with accession number GSE 108291, analyzed similarly as the original publication using the Seurat library (v4) (114, 115). Genes expressed in fewer than 3 cells along with cells with fewer than 200 detected genes were removed. Cells with more than 95% zero count genes, 4,700 mapped transcripts, and 7.5% mitochondrial gene expression were filtered out as well. Lastly, genes without annotations as well as mitochondrial and ribosomal genes were removed from the dataset for further analysis, which consisted of 6,037 cells and 15,747 genes. Using the SCTransform function, the dataset was simultaneously normalized, scaled, and regressed out factors related to biological doublet and cell cycle scored with the *CellCycleScoring* function. Clusters were segmented with 2,000 highly variable genes and the corresponding top 30 principal components at a 0.8 resolution, leading to 16 clusters. To assign identities of the clusters, differentially expressed genes were analyzed using a Wilcoxon rank sum test and compared to the marker genes in the original publication. However, we did not resolve two separate clusters for K6 S-Shaped Body and K7 Renal Vesicle; instead, our K7 Renal Vesicle/S-Shaped Body contained significantly upregulated genes for both populations. In our analysis, K10 and 15 had a very similar gene expression profile, both mapping well to the published K9 ureteric epithelium. Therefore, we grouped them together to achieve a total 15 unique clusters, similar to those reported by Combes and colleagues. To augment clustering resolution and retain consistency with the MATCHY clustering approach, we extracted the nephron lineage subpopulations (K0 primed nephron, K1 nephron progenitor, K5 CnS/distal tubule/Loop of Henle, K6 early proximal tubule, K7 renal vesicle/s-shaped body, and K12 committing NP/PTA) and reclustered them using the same marker gene set as in MATCHY. Two out of the 7 clusters lacked expression of MATCHY biomarkers but rather mapped to the proximal segment of the nephron. We therefore analyzed the remaining 5 clusters with the same differential gene expression mapping approach to identify N0 NPC, N2 PN, N5 PTA, N1 RV, and N3 BRV, as characterized in our biophysical measurements. Selected gene sets from the Broad Institute’s Molecular Signatures Database (MSigDB) were used to compare the expression levels between N0 NPC + N2 PN vs. N5 PTA + N1 RV to complete the Gene Set Enrichment Analysis (GSEA).

### Plotting and statistical analysis

One-way analysis of variance (ANOVA) with correction for multiple comparisons using Tukey’s honestly significant difference test was performed in MATLAB, along with paired and unpaired *t*-tests. Violin plots were generated in GraphPad Prism 10.

## References

1. C.-P. Heisenberg, Y. Bellaïche, Forces in tissue morphogenesis and patterning. Cell 153, 948–962 (2013).

2. S. Yang, et al., Morphogens enable interacting supracellular phases that generate organ architecture. Science 382, eadg5579 (2023).

3. A. E. Shyer, et al., Emergent cellular self-organization and mechanosensation initiate follicle pattern in the avian skin. Science 357, 811–815 (2017).

4. R. M. Houtekamer, M. C. van der Net, M. M. Maurice, M. Gloerich, Mechanical forces directing intestinal form and function. Curr. Biol. 32, R791–R805 (2022).

5. C. Parada, et al., Mechanical feedback defines organizing centers to drive digit emergence. Dev. Cell 57, 854–866.e6 (2022).

6. T. Mammoto, et al., Mechanochemical control of mesenchymal condensation and embryonic tooth organ formation. Dev. Cell 21, 758–769 (2011).

7. M. H. Dominguez, A. L. Krup, J. M. Muncie, B. G. Bruneau, Graded mesoderm assembly governs cell fate and morphogenesis of the early mammalian heart. Cell 186, 479–496.e23 (2023).

8. H. Tao, et al., Oscillatory cortical forces promote three dimensional cell intercalations that shape the murine mandibular arch. Nat. Commun. 10, 1703 (2019).

9. V. Osathanondh, E. L. Potter, Development of human kidney as shown by microdissection. IV. Development of tubular portions of nephrons. Arch. Pathol. 82, 391–402 (1966).

10. A. N. Combes, J. A. Davies, M. H. Little, Cell-cell interactions driving kidney morphogenesis. Curr. Top. Dev. Biol. 112, 467–508 (2015).

11. T. J. Carroll, J.-S. Park, S. Hayashi, A. Majumdar, A. P. McMahon, Wnt9b plays a central role in the regulation of mesenchymal to epithelial transitions underlying organogenesis of the mammalian urogenital system. Dev. Cell 9, 283–292 (2005).

12. J.-S. Park, M. T. Valerius, A. P. McMahon, Wnt/beta-catenin signaling regulates nephron induction during mouse kidney development. Development 134, 2533–2539 (2007).

13. A. C. Brown, et al., Role for compartmentalization in nephron progenitor differentiation. Proc. Natl. Acad. Sci. U. S. A. 110, 4640–4645 (2013).

14. A. Ihermann-Hella, et al., Dynamic MAPK/ERK Activity Sustains Nephron Progenitors through Niche Regulation and Primes Precursors for Differentiation. Stem Cell Reports 11, 912–928 (2018).

15. N. O. Lindström, N. O. Carragher, P. Hohenstein, The PI3K pathway balances self-renewal and differentiation of nephron progenitor cells through β-catenin signaling. Stem Cell Reports 4, 551– 560 (2015).

16. A. Reginensi, et al., Yap- and Cdc42-dependent nephrogenesis and morphogenesis during mouse kidney development. PLoS Genet. 9, e1003380 (2013).

17. A. Das, et al., Stromal-epithelial crosstalk regulates kidney progenitor cell differentiation. Nat. Cell Biol. 15, 1035–1044 (2013).

18. H. Wang, et al., STAT1 activation regulates proliferation and differentiation of renal progenitors. Cell. Signal. 22, 1717–1726 (2010).

19. N. O. Lindström, et al., Integrated β-catenin, BMP, PTEN, and Notch signalling patterns the nephron. Elife 3, e04000 (2015).

20. S. C. Boyle, M. Kim, M. T. Valerius, A. P. McMahon, R. Kopan, Notch pathway activation can replace the requirement for Wnt4 and Wnt9b in mesenchymal-to-epithelial transition of nephron stem cells. Development 138, 4245–4254 (2011).

21. E. Chung, P. Deacon, S. Marable, J. Shin, J.-S. Park, Notch signaling promotes nephrogenesis by downregulating Six2. Development 143, 3907–3913 (2016).

22. J. G. Lefevre, et al., Self-organisation after embryonic kidney dissociation is driven via selective adhesion of ureteric epithelial cells. Development 144, 1087–1096 (2017).

23. E. A. Cho, et al., Differential expression and function of cadherin-6 during renal epithelium development. Development 125, 803–812 (1998).

24. S. P. Mah, H. Saueressig, M. Goulding, C. Kintner, G. R. Dressler, Kidney development in cadherin-6 mutants: delayed mesenchyme-to-epithelial conversion and loss of nephrons. Dev. Biol. 223, 38–53 (2000).

25. S. Goto, et al., Involvement of R-cadherin in the early stage of glomerulogenesis. J. Am. Soc. Nephrol. 9, 1234–1241 (1998).

26. H. Y. Kim, T. R. Jackson, L. A. Davidson, On the role of mechanics in driving mesenchymal-to-epithelial transitions. Semin. Cell Dev. Biol. 67, 113–122 (2017).

27. A. Gupta, M. P. Lutolf, A. J. Hughes, K. F. Sonnen, Bioengineering in vitro models of embryonic development. Stem Cell Reports 16, 1104–1116 (2021).

28. G. Binnig, C. F. Quate, C. Gerber, Atomic force microscope. Phys. Rev. Lett. 56, 930–933 (1986).

29. H.-J. Butt, B. Cappella, M. Kappl, Force measurements with the atomic force microscope: Technique, interpretation and applications. Surf. Sci. Rep. 59, 1–152 (2005).

30. B. A. Nerger, et al., 3D Hydrogel Encapsulation Regulates Nephrogenesis in Kidney Organoids. Adv. Mater., e2308325 (2024).

31. R. M. Hochmuth, Micropipette aspiration of living cells. J. Biomech. 33, 15–22 (2000).

32. B. Hogan, A. Babataheri, Y. Hwang, A. I. Barakat, J. Husson, Characterizing cell adhesion by using micropipette aspiration. Biophys. J. 109, 209–219 (2015).

33. T. Y.-C. Tsai, et al., An adhesion code ensures robust pattern formation during tissue morphogenesis. Science 370, 113–116 (2020).

34. H. Zhang, K.-K. Liu, Optical tweezers for single cells. J. R. Soc. Interface 5, 671–690 (2008).

35. L. Novotny, R. X. Bian, X. S. Xie, Theory of Nanometric Optical Tweezers. Phys. Rev. Lett. 79, 645–648 (1997).

36. D. Molino, et al., On-Chip Quantitative Measurement of Mechanical Stresses During Cell Migration with Emulsion Droplets. Sci. Rep. 6, 29113 (2016).

37. F. Serwane, et al., In vivo quantification of spatially varying mechanical properties in developing tissues. Nat. Methods 14, 181–186 (2017).

38. M. Dembo, Y.-L. Wang, Stresses at the Cell-to-Substrate Interface during Locomotion of Fibroblasts. Biophys. J. 76, 2307–2316 (1999).

39. B. Sabass, M. L. Gardel, C. M. Waterman, U. S. Schwarz, High resolution traction force microscopy based on experimental and computational advances. Biophys. J. 94, 207–220 (2008).

40. M. Bao, et al., Stem cell-derived synthetic embryos self-assemble by exploiting cadherin codes and cortical tension. Nat. Cell Biol. 24, 1341–1349 (2022).

41. M. Lusis, et al., Isolation of clonogenic, long-term self renewing embryonic renal stem cells. Stem Cell Res. 5, 23–39 (2010).

42. C. Xinaris, et al., In vivo maturation of functional renal organoids formed from embryonic cell suspensions. J. Am. Soc. Nephrol. 23, 1857–1868 (2012).

43. A. E. Cerchiari, et al., A strategy for tissue self-organization that is robust to cellular heterogeneity and plasticity. Proc. Natl. Acad. Sci. U. S. A. 112, 2287–2292 (2015).

44. M. C. Recuenco, et al., Nonmuscle myosin II regulates the morphogenesis of metanephric mesenchyme-derived immature nephrons. J. Am. Soc. Nephrol. 26, 1081–1091 (2015).

45. U. Muller, et al., Integrin ɑ8β1 Is Critically Important for Epithelial–Mesenchymal Interactions during Kidney Morphogenesis. Cell 88, 603–613 (1997).

46. K. Leclerc, F. Costantini, Mosaic analysis of cell rearrangements during ureteric bud branching in dissociated/reaggregated kidney cultures and in vivo. Dev. Dyn. 245, 483–496 (2016).

47. M. Unbekandt, J. A. Davies, Dissociation of embryonic kidneys followed by reaggregation allows the formation of renal tissues. Kidney Int. 77, 407–416 (2010).

48. V. Ganeva, M. Unbekandt, J. A. Davies, An improved kidney dissociation and reaggregation culture system results in nephrons arranged organotypically around a single collecting duct system. Organogenesis 7, 83–87 (2011).

49. B. Der, H. Bugacov, B.-M. Briantseva, A. P. McMahon, Cadherin Adhesion Complexes Direct Cell Aggregation in the Epithelial Transition of Wnt-Induced Nephron Progenitor Cells. *bioRxiv*, 2023.08.27.555021 (2023).

50. U. Dahl, et al., Genetic dissection of cadherin function during nephrogenesis. Mol. Cell. Biol. 22, 1474–1487 (2002).

51. T. Y.-C. Tsai, R. M. Garner, S. G. Megason, Adhesion-Based Self-Organization in Tissue Patterning. Annu. Rev. Cell Dev. Biol. 38, 349–374 (2022).

52. M. S. Steinberg, Reconstruction of Tissues by Dissociated Cells. Science 141, 401–408 (1963).

53. G. W. Brodland, The Differential Interfacial Tension Hypothesis (DITH): A Comprehensive Theory for the Self-Rearrangement of Embryonic Cells and Tissues. J. Biomech. Eng. 124, 188–197 (2002).

54. L. Saxen, J. Wartiovaara, Cell contact and cell adhesion during tissue organization. Int. J. Cancer 1, 271–290 (1966).

55. M. J. Tang, D. Worley, M. Sanicola, G. R. Dressler, The RET-glial cell-derived neurotrophic factor (GDNF) pathway stimulates migration and chemoattraction of epithelial cells. J. Cell Biol. 142, 1337–1345 (1998).

56. C. E. Fisher, L. Michael, M. W. Barnett, J. A. Davies, Erk MAP kinase regulates branching morphogenesis in the developing mouse kidney. Development 128, 4329–4338 (2001).

57. N. Hino, et al., ERK-Mediated Mechanochemical Waves Direct Collective Cell Polarization. Dev. Cell 53, 646–660.e8 (2020).

58. J. N. Lakins, A. R. Chin, V. M. Weaver, Exploring the link between human embryonic stem cell organization and fate using tension-calibrated extracellular matrix functionalized polyacrylamide gels. Methods Mol. Biol. 916, 317–350 (2012).

59. T. Vignaud, et al., Stress fibres are embedded in a contractile cortical network. Nat. Mater. 20, 410–420 (2021).

60. Q. Tseng, et al., Spatial organization of the extracellular matrix regulates cell–cell junction positioning. Proceedings of the National Academy of Sciences 109, 1506–1511 (2012).

61. J. M. Viola, et al., Rho/ROCK activity tunes cell compartment segregation and differentiation in nephron-forming niches. bioRxiv, 2023.11.08.566308 (2023).

62. A. Schuchardt, V. D’Agati, L. Larsson-Blomberg, F. Costantini, V. Pachnis, Defects in the kidney and enteric nervous system of mice lacking the tyrosine kinase receptor Ret. Nature 367, 380–383 (1994).

63. G. Cacalano, et al., GFRa1 Is an Essential Receptor Component for GDNF in the Developing Nervous System and Kidney. Neuron 21, 53–62 (1998).

64. L. Michael, D. E. Sweeney, J. A. Davies, A role for microfilament-based contraction in branching morphogenesis of the ureteric bud. Kidney Int. 68, 2010–2018 (2005).

65. M. V. Barone, et al., RET/PTC1 oncogene signaling in PC Cl 3 thyroid cells requires the small GTP-binding protein Rho. Oncogene 20, 6973–6982 (2001).

66. L. Saxén, H. Sariola, Early organogenesis of the kidney. Pediatr. Nephrol. 1, 385–392 (1987).

67. K. M. Short, et al., Global quantification of tissue dynamics in the developing mouse kidney. Dev. Cell 29, 188–202 (2014).

68. V. Osathanondh, E. L. Potter, DEVELOPMENT OF HUMAN KIDNEY AS SHOWN BY MICRODISSECTION. III. FORMATION AND INTERRELATIONSHIP OF COLLECTING TUBULES AND NEPHRONS. Arch. Pathol. 76, 290–302 (1963).

69. K. Stark, S. Vainio, G. Vassileva, A. P. McMahon, Epithelial transformation of metanephric mesenchyme in the developing kidney regulated by Wnt-4. Nature 372, 679–683 (1994).

70. L. Canty, E. Zarour, L. Kashkooli, P. François, F. Fagotto, Sorting at embryonic boundaries requires high heterotypic interfacial tension. Nat. Commun. 8, 157 (2017).

71. M. L. Manning, R. A. Foty, M. S. Steinberg, E.-M. Schoetz, Coaction of intercellular adhesion and cortical tension specifies tissue surface tension. Proc. Natl. Acad. Sci. U. S. A. 107, 12517–12522 (2010).

72. G. Klein, M. Langegger, C. Goridis, P. Ekblom, Neural cell adhesion molecules during embryonic induction and development of the kidney. Development 102, 749–761 (1988).

73. Y. Kimura, et al., Cadherin-11 expressed in association with mesenchymal morphogenesis in the head, somite, and limb bud of early mouse embryos. Dev. Biol. 169, 347–358 (1995).

74. D. Vestweber, R. Kemler, P. Ekblom, Cell-adhesion molecule uvomorulin during kidney development. Dev. Biol. 112, 213–221 (1985).

75. J. A. Davies, D. R. Garrod, Induction of early stages of kidney tubule differentiation by lithium ions. Dev. Biol. 167, 50–60 (1995).

76. A. N. Combes, et al., Single cell analysis of the developing mouse kidney provides deeper insight into marker gene expression and ligand-receptor crosstalk. Development 146, dev178673 (2019).

77. K. Kretzschmar, F. M. Watt, Lineage tracing. Cell 148, 33–45 (2012).

78. K. Georgas, et al., Analysis of early nephron patterning reveals a role for distal RV proliferation in fusion to the ureteric tip via a cap mesenchyme-derived connecting segment. Dev. Biol. 332, 273– 286 (2009).

79. N. O. Lindström, et al., Progressive Recruitment of Mesenchymal Progenitors Reveals a Time-Dependent Process of Cell Fate Acquisition in Mouse and Human Nephrogenesis. Dev. Cell 45, 651–660.e4 (2018).

80. N. O. Lindström, et al., Spatial transcriptional mapping of the human nephrogenic program. Dev. Cell 56, 2381–2398.e6 (2021).

81. P. Tikka, et al., Computational modelling of nephron progenitor cell movement and aggregation during kidney organogenesis. Math. Biosci. 344, 108759 (2022).

82. A. N. Combes, J. G. Lefevre, S. Wilson, N. A. Hamilton, M. H. Little, Cap mesenchyme cell swarming during kidney development is influenced by attraction, repulsion, and adhesion to the ureteric tip. Dev. Biol. 418, 297–306 (2016).

83. L. L. O’Brien, et al., Wnt11 directs nephron progenitor polarity and motile behavior ultimately determining nephron endowment. Elife 7 (2018).

84. N. Naiman, et al., Repression of Interstitial Identity in Nephron Progenitor Cells by Pax2 Establishes the Nephron-Interstitium Boundary during Kidney Development. Dev. Cell 41, 349–365.e3 (2017).

85. C. M. Karner, et al., Wnt9b signaling regulates planar cell polarity and kidney tubule morphogenesis. Nat. Genet. 41, 793–799 (2009).

86. M. H. Little, A. N. Combes, Kidney organoids: accurate models or fortunate accidents. Genes Dev. 33, 1319–1345 (2019).

87. R. Kopan, H.-T. Cheng, K. Surendran, Molecular insights into segmentation along the proximal-distal axis of the nephron. J. Am. Soc. Nephrol. 18, 2014–2020 (2007).

88. C. R. Pfeifer, A. E. Shyer, A. R. Rodrigues, Creative processes during vertebrate organ morphogenesis: Biophysical self-organization at the supracellular scale. Curr. Opin. Cell Biol. 86, 102305 (2024).

89. M. Mederacke, L. Conrad, N. Doumpas, R. Vetter, D. Iber, Geometric effects position renal vesicles during kidney development. Cell Rep. 42, 113526 (2023).

90. H. Ramalingam, et al., Disparate levels of beta-catenin activity determine nephron progenitor cell fate. Dev. Biol. 440, 13–21 (2018).

91. A. R. England, et al., Identification and characterization of cellular heterogeneity within the developing renal interstitium. Development 147 (2020).

92. S. B. Wilson, M. H. Little, The origin and role of the renal stroma. Development 148 (2021).

93. O. Campàs, A toolbox to explore the mechanics of living embryonic tissues. Semin. Cell Dev. Biol. 55, 119–130 (2016).

94. N. Yamaguchi, et al., Rear traction forces drive adherent tissue migration in vivo. Nat. Cell Biol. 24, 194–204 (2022).

95. A. Mongera, et al., A fluid-to-solid jamming transition underlies vertebrate body axis elongation. Nature 561, 401–405 (2018).

96. J. M. Viola, et al., Tubule jamming in the developing mouse kidney creates cyclical mechanical stresses in nephron-forming niches. bioRxiv, 2022.06.03.494718 (2023).

97. G. Scarcelli, et al., Noncontact three-dimensional mapping of intracellular hydromechanical properties by Brillouin microscopy. Nat. Methods 12, 1132–1134 (2015).

98. J. Zhang, G. Scarcelli, Mapping mechanical properties of biological materials via an add-on Brillouin module to confocal microscopes. Nat. Protoc. 16, 1251–1275 (2021).

99. K. Sugimura, P.-F. Lenne, F. Graner, Measuring forces and stresses in situ in living tissues. Development 143, 186–196 (2016).

100. A. C. Daly, M. E. Prendergast, A. J. Hughes, J. A. Burdick, Bioprinting for the Biologist. Cell 184, 18–32 (2021).

101. S. E. Howden, et al., Plasticity of distal nephron epithelia from human kidney organoids enables the induction of ureteric tip and stalk. Cell Stem Cell 28, 671–684.e6 (2021).

102. S. V. Kumar, et al., Kidney micro-organoids in suspension culture as a scalable source of human pluripotent stem cell-derived kidney cells. Development 146 (2019).

103. N. Wanek, K. Muneoka, G. Holler-Dinsmore, R. Burton, S. V. Bryant, A staging system for mouse limb development. J. Exp. Zool. 249, 41–49 (1989).

104. H. Barak, S. C. Boyle, Organ culture and immunostaining of mouse embryonic kidneys. Cold Spring Harb. Protoc. 2011, db.prot5558 (2011).

105. L. S. Prahl, J. M. Viola, J. Liu, A. J. Hughes, The developing kidney actively negotiates geometric packing conflicts to avoid defects. bioRxiv, 2021.11.29.470441 (2021).

106. A. N. Combes, et al., An integrated pipeline for the multidimensional analysis of branching morphogenesis. Nat. Protoc. 9, 2859–2879 (2014).

107. A. C. Brown, et al., Isolation and culture of cells from the nephrogenic zone of the embryonic mouse kidney. J. Vis. Exp. (2011) 10.3791/2555.

108. A. C. Brown, S. D. Muthukrishnan, L. Oxburgh, A synthetic niche for nephron progenitor cells. Dev. Cell 34, 229–241 (2015).

109. J. M. Viola, et al., Guiding cell network assembly using shape-morphing hydrogels. Adv. Mater. 32, e2002195 (2020).

110. L. S. Prahl, C. M. Porter, J. Liu, J. M. Viola, A. J. Hughes, Independent control over cell patterning and adhesion on hydrogel substrates for tissue interface mechanobiology. iScience 26, 106657 (2023).

111. J. Schindelin, et al., Fiji: an open-source platform for biological-image analysis. Nat. Methods 9, 676–682 (2012).

112. D. G. Lowe, Distinctive Image Features from Scale-Invariant Keypoints. International Journal of Computer Vision 60, 91–110 (2004).

113. U. Schmidt, M. Weigert, C. Broaddus, G. Myers, Cell Detection with Star-Convex Polygons in Medical Image Computing and Computer Assisted Intervention – MICCAI 2018, (Springer International Publishing, 2018), pp. 265–273.

114. R. Satija, J. A. Farrell, D. Gennert, A. F. Schier, A. Regev, Spatial reconstruction of single-cell gene expression data. Nat. Biotechnol. 33, 495–502 (2015).

115. A. Butler, P. Hoffman, P. Smibert, E. Papalexi, R. Satija, Integrating single-cell transcriptomic data across different conditions, technologies, and species. Nat. Biotechnol. 36, 411–420 (2018).

